# Group Based Trajectory Analysis of Cognitive Outcomes in Children with Perinatal HIV

**DOI:** 10.1101/398339

**Authors:** Payal B. Patel, Tanakorn Apornpong, Stephen J. Kerr, Thanyawee Puthanakit, K. Thongpibul, P. Kosalaraksa, P. Ounchanum, S. Kanjanavanit, C. Ngampiyaskul, W. Luesomboon, L. Penhusun, K. Chettra, Claude Mellins, Kay Malee, Serena Spudich, Jintanat Ananworanich, Robert Paul, On behalf of the PREDICT/Resilience Study Group

**Affiliations:** Department of Neurology, Yale University; HIV-NAT, The Thai Red Cross AIDS Research Center, Bangkok, Thailand; Department of Pediatrics, Faculty of Medicine, Chulalongkorn University, Bangkok, Thailand; Faculty of Medicine, Chiang Mai University, Bangkok, Thailand; Department of Pediatrics, Faculty of Medicine, Khon Kaen University, Khon Kaen, Thailand; Chiangrai Prachanukroh Hospital, Chiang Rai, Thailand; Nakornping Hospital, Chiang Mai, Thailand; Prapokklao Hospital, Chantaburi, Thailand; Queen Savang Vadhana Memorial Hospital, Chonburi, Thailand; National Center for HIV/AIDS Dermatology and STDs, Phnom Penh, Cambodia; HIV Center for Clinical and Behavioral Studies, New York State Psychiatric Institute, and Columbia University, New York, USA; Northwestern University Feinberg School of Medicine, Department of Psychiatry and Behavioral Science, Chicago IL; SEARCH, Thai Red Cross AIDS Research Center, Bangkok, Thailand; The Henry M. Jackson Foundation for the Advancement of Military Medicine, Bethesda, MD, USA; Department of Global Health, University of Amsterdam, Amsterdam, Netherlands; Missouri Institute of Mental Health, University of Missouri-St. Louis, MO, USA

## Abstract

**Background:** Cognitive impairment is common in children with perinatally-acquired HIV (pHIV). It is not known whether exposure to HIV-related neuropathogenic mechanisms during vulnerable periods of neurodevelopment may produce distinct long-term cognitive phenotypes as children age. We used group based trajectory modeling to identify clusters of children with pHIV following a unique developmental trajectory across age and predictors of belonging to select cognitive trajectory groups.

**Methods:** Participants included children aged 1 to 17 enrolled in the PREDICT resilience study, a cohort study of children with pHIV in Thailand and Cambodia. Cognitive testing was conducted semi-annually over three years. Group based trajectory analyses determined subgroups of children with differing cognitive trajectories using maximum likelihood estimates and Bayesian statistics. Multiple logistic regression identified baseline factors associated with belonging to the lowest scoring trajectory group.

**Results:** Three distinct cognitive phenotypes were identified for each neurocognitive test categorized as high, medium and low scoring groups. A subgroup of children demonstrated normal developmental patterns for Color Trails Test 1 and 2. Children in the low trajectory group were more likely to present at an older age (>8 years, OR: 2.72; p 0.01) and report lower household income level (OR: 0.33-0.42; p<0.005). Neither CD4 nadir nor treatment arm was associated with cognitive trajectory status.

**Conclusion:** Our study reflects the benefit of using group based trajectory modeling to classify the heterogeneity in cognitive outcomes of children with pHIV. Children were described as belonging to three distinct subgroups determined at study onset alluding to the fact that cognitive outcomes are likely to be determined at an early age with little variability over time in children with pHIV. Demographic variables, including older age at presentation and household income, were associated with low scoring cognitive trajectories, whereas HIV related variables were not. These findings mirror other studies and demonstrate the impact of socioeconomic factors on cognitive development in children with pHIV.

## Introduction

Cognitive impairment is common in children with perinatal HIV (pHIV) even in those who are virally suppressed on anti-retroviral therapy (ART)^1^. Working memory, processing speed and executive function domains are primarily affected in children living with HIV^2^. While the cross-sectional profile of cognitive impairment in children with HIV has been well described^3^, less is known about the developmental trajectory of children and adolescents living with pHIV^4^. Of particular interest is whether subgroups of children with pHIV can be classified based on longitudinal cognitive performance. Additionally, definitions of cognitive impairment vary between studies, limiting our ability to assess prevalence of cognitive impairment in children living with HIV globally, especially in cohorts residing in resource-limited settings.

Group-based trajectory modeling addresses these issues by identifying clusters of individuals that follow consistent paths over time without ex ante criteria for cognitive impairment^5^. Longitudinal data can be summarized in a visually transparent fashion and subgroups of children with differing developmental trajectories can be compared in the absence of normative data using group based trajectory modeling.

Our previous work conducted utilized multivariate linear regression analyses to compare cognitive scores at select time points in our pediatric HIV cohort in Thailand and Cambodia revealed significantly worse cognitive performance in the pHIV group compared to control participants at baseline and again at 144 weeks regardless of treatment history^6^. Here, we expand on these findings by employing group based trajectory modeling to identify subgroups of children with distinct cognitive profiles. Baseline predictors of low scoring trajectory subgroups were examined. Identification of early factors associated with poor cognitive outcomes in children with perinatally acquired HIV (pHIV) may provide the basis for early screening and intervention in this vulnerable population.

## Study Design

Participants included children enrolled in the Pediatric Randomized Early versus Deferred Initiation in Cambodia and Thailand (PREDICT) study at 7 sites across Thailand and 2 sites in Cambodia^7^. ART-naïve children between the ages of 1-12 years with CD4% between 15-24% and no history of AIDS defining illness were included in the study. Information regarding sex, age, ethnicity, CD4 nadir, caregiver status and household income was collected. Caregiver status was classified as living with biological parent, living with other relative, or living in orphanage. Household income was determined by caregiver self-report and stratified as below average, average or above average based on national data. Laboratory and clinical evaluations were completed every 12 weeks and included general history, CD4 count and percentage. Viral load was measured every 24 weeks. Children were randomized to initiate ART at according to the national guidelines operative at the time (CD4 < 15% or CDC category C event) or earlier (CD4 < 25%). First line ART regimen included zidovudine, lamivudine and nevirapine. In children with prior exposure to nevirapine as part of prophylaxis for maternal to child transmission, a protease inhibitor was used as a substitute. Informed consent from caregivers was obtained in writing prior to study enrollment. The study was approved by Thai and Cambodian national and local Institutional Review Boards. After completion of the PREDICT study (ISRCTN00234091), most children elected to enroll in a subsequent longitudinal study (Resilience; (MH102151). Data included in this study were extracted from both PREDICT and Resilience.

### Neuropsychological testing

Neuropsychological testing was completed annually by Thai psychologists or trained nurses as a sub-study of the main PREDICT trial beginning in 2008. Trained nurses were certified after correctly completing and scoring a minimum of 10 participants per test under supervision. Testing included translated versions of Wechsler Intelligence Scale for Children (WISC-III) ages 6 years and older or Wechsler Preschool and Primary Scale of Intelligence (WPPSI) for ages 2 to 5, Beery Visual Motor Integration Scale (Beery VMI), and Color Trails Test 1 and 2 (CTT1 and CTT2). Previously translated and validated Thai versions of WISC-III and WPPSI were used for Thai children only. Berry VMI and CTT instructions were translated into Thai and Khmer languages by bi-lingual translators^8^. External quality assurance was performed by randomly selected video recording, observation and scoring review by a US based neuropsychologist. Raw scores for all tests were transformed to scaled scores using US-based norms.

### Statistical analyses

A modified approach to group based trajectory analysis (GBTA) was conducted to identify subgroups of children following similar cognitive trajectories across age (Nagin, 2005). Neuropsychological tests were analyzed independently. Cognitive scores for each participant across the 144 weeks of study were categorized as a function of age using maximum likelihood estimation as described by Nagin et al^5^, adjusting for the time varying covariates of log viral load and treatment status (on or off ART). Joint estimation of the parameters describing trajectory shape and time varying covariates allowed us to account for the influence of variable treatment initiation on trajectory group membership. Best fit of trajectory group number and shape was determined using Bayesian Information Criterion^9^. Individuals were assigned to the group for which the posterior probability of membership was highest. Posterior probabilities and odds ratios were used to assess the adequacy of our model. Second, we assessed change in cognitive test scores as participants aged using longitudinal random effects linear models and the lowest age at which the test was conducted as a reference group. Third, multiple logistic regression was used to identify factors associated with belonging to the lowest scoring trajectory strata for each cognitive test. Variables in univariate analyses achieving threshold of p<0.15 were retained in multivariate models.

## Results

Among the 286 children enrolled, all completed the Beery Visuomotor Integration Scale (Beery VMI), 264 (92%) completed the Color Trails Tests (CTT) 1 and 2, and 165 (58%) completed IQ testing. Demographic and treatment information are described in Table 1. Over half of children were on ART at the start of the neurocognitive sub-study and the majority (87%) were on treatment at the end of the study.

**Table 1.**
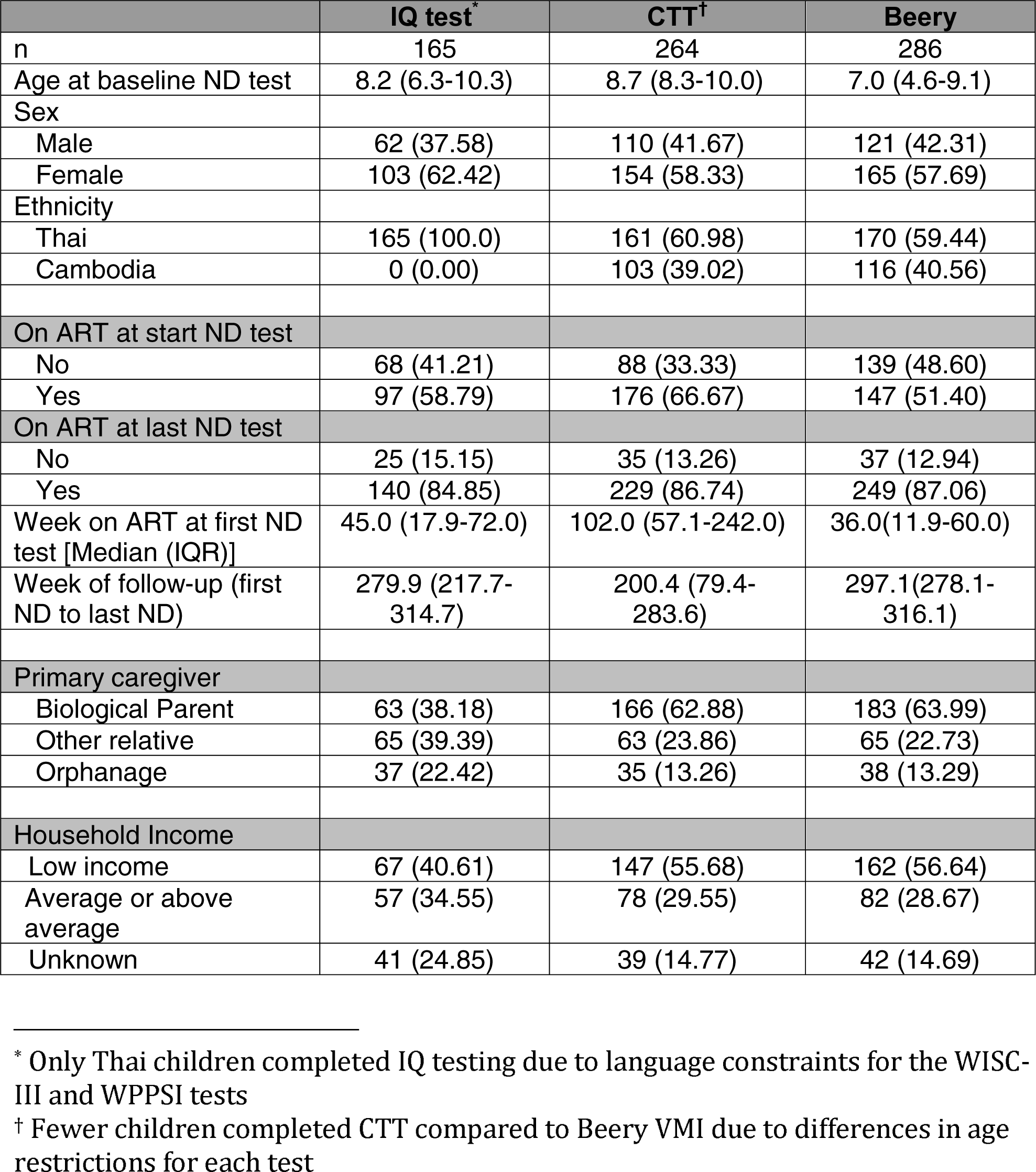
Demographic and treatment variables

|  | **IQ test\*** | **CTT†** | **Beery** |
| --- | --- | --- | --- |
| n | 165 | 264 | 286 |
| Age at baseline ND test | 8.2 (6.3-10.3) | 8.7 (8.3-10.0) | 7.0 (4.6-9.1) |
| Sex |  |  |  |
| Male | 62 (37.58) | 110 (41.67) | 121 (42.31) |
| Female | 103 (62.42) | 154 (58.33) | 165 (57.69) |
| Ethnicity |  |  |  |
| Thai | 165 (100.0) | 161 (60.98) | 170 (59.44) |
| Cambodia | 0 (0.00) | 103 (39.02) | 116 (40.56) |
| On ART at start ND test |  |  |  |
| No | 68 (41.21) | 88 (33.33) | 139 (48.60) |
| Yes | 97 (58.79) | 176 (66.67) | 147 (51.40) |
| On ART at last ND test |  |  |  |
| No | 25 (15.15) | 35 (13.26) | 37 (12.94) |
| Yes | 140 (84.85) | 229 (86.74) | 249 (87.06) |
| Week on ART at first ND test [Median (IQR)] | 45.0 (17.9-72.0) | 102.0 (57.1-242.0) | 36.0(11.9-60.0) |
| Week of follow-up (first ND to last ND) | 279.9 (217.7-314.7) | 200.4 (79.4-283.6) | 297.1(278.1-316.1) |
| Primary caregiver |  |  |  |
| Biological Parent | 63 (38.18) | 166 (62.88) | 183 (63.99) |
| Other relative | 65 (39.39) | 63 (23.86) | 65 (22.73) |
| Orphanage | 37 (22.42) | 35 (13.26) | 38 (13.29) |
| Household Income |  |  |  |
| Low income | 67 (40.61) | 147 (55.68) | 162 (56.64) |
| Average or above average | 57 (34.55) | 78 (29.55) | 82 (28.67) |
| Unknown | 41 (24.85) | 39 (14.77) | 42 (14.69) |

\* Only Thai children completed IQ testing due to language constraints for the WISC-III and WPPSI tests
† Fewer children completed CTT compared to Beery VMI due to differences in age restrictions for each test

Graphs 1-3 show trajectory models for each cognitive test after adjusting for time-varying covariates of treatment status and viral load. To assess model fit, posterior probabilities for individual group assignments were ≥ 0.7 and odds for individual group assignment were ≥ 4.5 in all trajectory groups. For each neurocognitive test, we identified three unique trajectory groups, classified as high, medium and low performance groups. We identified similar trends across all trajectory groups for select cognitive tests. Verbal IQ scores decreased for each trajectory by approximately 10 points across childhood and into early adolescence. Beery VMI scores and Performance IQ scores remained relatively stable across age. In contrast, CTT 1 & 2 scores in the high scoring trajectory group showed a 10-increase across older childhood and early adolescence, while the low and medium scoring trajectories showed no change across age. Numbers and mean scores across age for each trajectory group are provided (supplement table).

Results of multivariate logistic regression models identifying risk factors predictive of low cognitive group status are presented in Table 2. Average or above average household income was associated with reduced odds of belonging to the lowest cognitive trajectory groups across multiple cognitive tests (OR 0.33-0.42; p<0.005). For Beery VMI, children not living with their biological parent had reduced odds of belonging to the lowest scoring trajectory group (OR: 0.42, p=0.01). For Performance IQ, older age (> 8 years) at time of presentation was associated with increased odds of low cognitive group status (OR 2.72; p=0.01). HIV-related variables of CD4 nadir (CD4 count <350) and treatment arm (immediate ART vs deferred ART) were not associated with belonging to a select trajectory group.

**Table 2.** Significant baseline risk factors associated with low cognitive trajectory group status in multivariate analyses

| Performance IQ | OR | 95%CI | P |
| --- | --- | --- | --- |
| **Age at first entry to trajectory** |  |  |  |
| <8 | Ref |  |  |
| >8 | 2.72 | (1.31-5.65) | 0.01 |
| **Color Trails Test 1** | **OR** | **95%CI** | **P** |
| **Household Income** |  |  |  |
| Low income | Ref |  |  |
| Average or above average | 0.33 | (0.17-0.65) | 0.001 |
| Unknown | 0.87 | (0.32-2.35) | 0.79 |
| **Berry VMI** | **OR** | **95%CI** | **P** |
| **Household Income** |  |  |  |
| Low income | Ref |  |  |
| Average or above average | 0.41 | (0.22-0.74) | 0.003 |
| Unknown | 2.56 | (0.95-6.94) | 0.06 |
| **Primary caregiver** |  |  |  |
| Parent | Ref |  |  |
| Other | 0.42 | (0.22-0.82) | 0.01 |

## Discussion

Group based trajectory analyses displayed significant heterogeneity in cognitive outcomes among our cohort of Southeast Asian children who are long-term survivors of perinatal HIV. These results expand on our prior analyses, which focused on mean standardized scores at select time intervals, and allowed us to identify subgroups of children with pHIV with relatively poor cognitive outcomes across all age groups. Our study reflects the benefits of utilizing this novel approach, which is not reliant on strict clinical classifications to analyze cognitive outcomes. Criteria for neurocognitive impairment vary between studies and lack of local or appropriately controlled normative data can result in inaccurate representations of the prevalence of cognitive impairment in children with pHIV living in resource-limited settings^10^. By calibrating our model to calculate classification in terms of probability or odds of cognitive group membership, we were able to summarize complex longitudinal outcomes across an age spectrum and assess for risk factors associated with poor outcomes in a succinct and transparent manner.

Decline in verbal IQ scores in all trajectory groups as children age is most likely representative of limitations related to applying neurocognitive tests developed for native English speakers to non-English speaking children and adolescents in resource-limited settings. The increase in scores across age in the high scoring trajectory group for CTT 1 & 2 reflect children with normal developmental patterns for motor and executive function domains as CTT scores tend to increase as children age^11^, while the low and medium scoring groups with stagnant scores are potentially representative of a subgroup of children who are not achieving developmental gains due to low cognitive reserve. These results demonstrate the utility of group based trajectory analyses in identifying subgroups of children with diverse cognitive outcomes across the age spectrum.

Prior work has demonstrated the strong impact of socioeconomic factors on cognitive outcomes in children with and without HIV^12,13^. Our results indicate that certain environmental factors, poverty in particular, are associated with poor cognitive outcomes independent of disease severity and treatment in children who are long-term survivors of HIV. Poverty is a known indicator for poor developmental outcomes in children residing in low and middle-income countries, including Thailand and Cambodia^14,15^. A study on adolescents and young adults living with HIV in the UK demonstrated strong correspondence between poverty and neurocognitive scores, while HIV-related factors in youth without prior AIDS defining illness were not associated with cognitive outcomes^3^ mirroring findings in our study that socioeconomic factors play a significant role in cognitive development independent of HIV severity in children without prior AIDS defining illnesses.

Interesting, not having a biologic parent as the primary caregiver was not associated with low cognitive group status on the Berry VMI. Studies performed in Thailand have shown that a large portion of children living with HIV are cared for by a relative who is not their biological parent as was found in our cohort^16^. Further, cognitive outcomes in pHIV are typically associated with better cognitive outcomes^16^. Caregiver stress negatively influences cognitive and behavioral functioning for children living with HIV^17^. These studies underscore the importance of incorporating family and caregiver mental health as part of the continuum of care for pHIV. Additional studies are needed to determine if the association between caregiver status and Beery VMI is mediated by overlapping psychosocial factors.

Our earlier publication demonstrated that differential initiation of ART (i.e. at presentation or following immunosuppression) in children with HIV presenting after infancy did not influence neurodevelopmental outcomes^6^. The current study expands on these findings and found that older age (>8 years) at study enrollment was associated with lower performance on IQ scores. This result demonstrates that, although timing of ART initiation did not influence cognitive status, later presentation to care is a possible risk factor for poor cognitive outcomes in children with pHIV. Prior studies evaluating ART initiation in children with HIV suggest a varied response that is dependent on an individual’s age at presentation^18,19^. PHIV+ infants and young children who initiated ART and were virally suppressed prior to 5 years of age had improved IQ scores in older childhood^20,21^, whereas children who began ART at an older age did not benefit from immediate initiation of antiretroviral therapy^4,6^. These results along with our findings suggest that initiation of ART in infancy or early childhood may be protective against poor cognitive outcomes.

Our results should be applied with caution as we evaluated cognitive trajectories in children at various ages over a select interval of time, and a limited number of children in our study were enrolled in early childhood (<5 years) and older adolescence (>14 years). Additionally, the clinical relevance of these results is difficult to assess without expected normative trajectories for Thai and Cambodian children. There are likely many factors associated with cognitive outcomes that were not included in our study, including effect modifiers of the association between poverty and cognitive outcomes. Regardless, our study found that developmental outcomes in children with pHIV are comprised of unique cognitive profiles across age as demonstrated through the use of group based trajectory modeling. Longer follow-up including transition into older adolescence and assessment of HIV-exposed, but uninfected children will be the focus of future studies.

---

**Graph 1:**
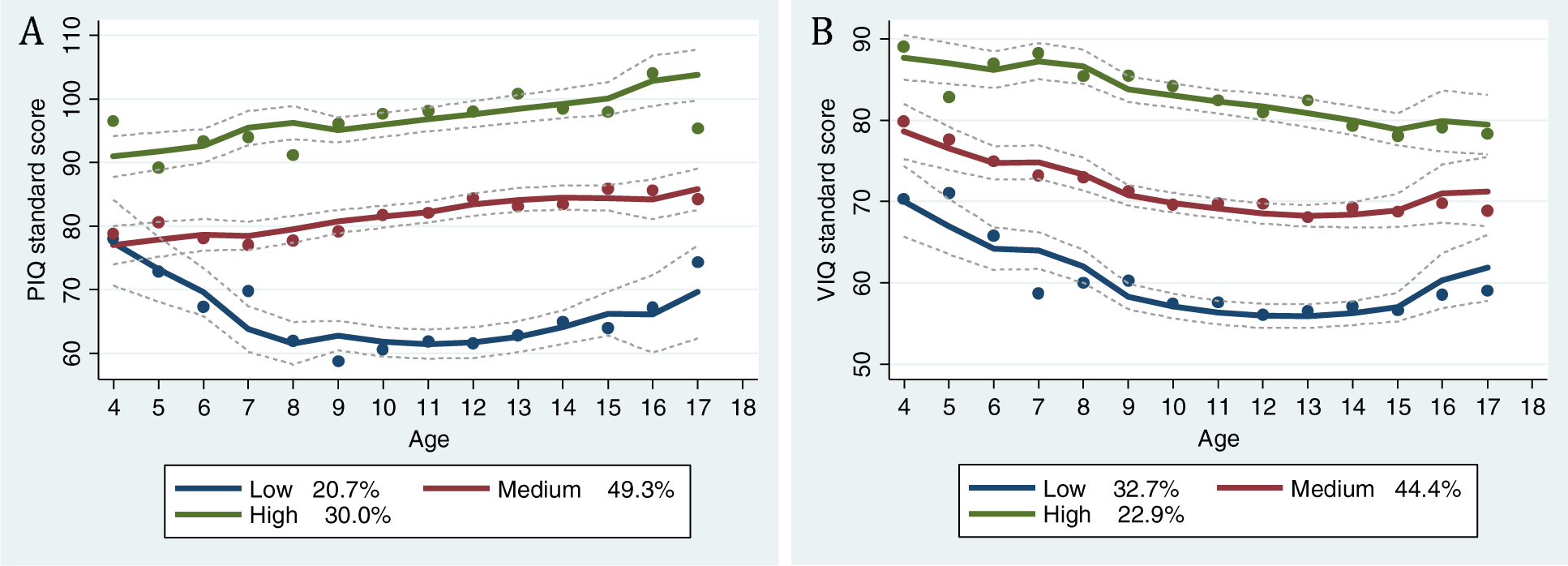
IQ trajectory groups. A. This graph demonstrates three trajectory subgroups, characterized as high, medium and low, for performance IQ test and percentage of children of children belonging to each subgroup. Scores in each subgroup remain relatively stable across age. B. This graph demonstrates three trajectory subgroups, characterized as high, medium and low, for verbal IQ test and percentage of children belonging to each subgroup. Scores in all groups decreased across age by approximately 10 points.

**Graph 2:**
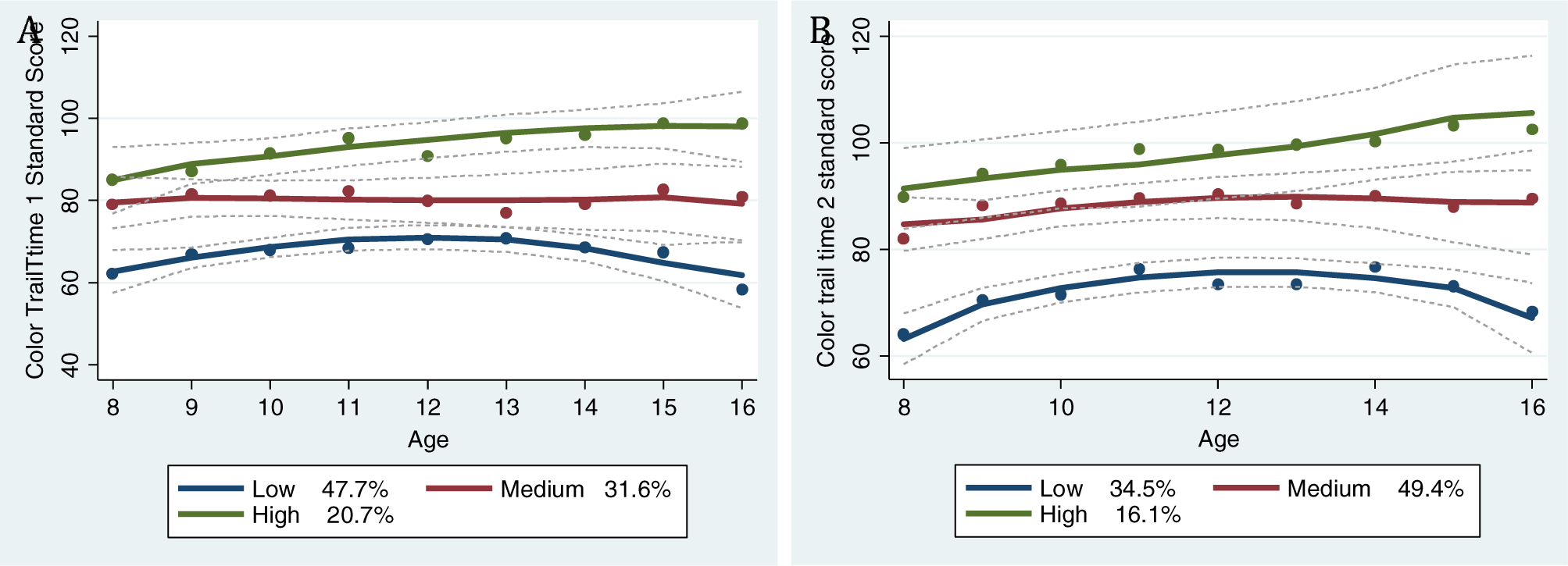
Color Trail Tests trajectory group. A. This graph demonstrates three trajectory subgroups, characterized as high, medium and low, for Color Trails Test 1 and percentage of children of children belonging to each subgroup. Scores in the low and medium trajectory groups remained stable, while scores in high group increased by 10 pt across age. B. This graph demonstrates three trajectory subgroups, characterized as high, medium and low, for Color Trails Test 2 and percentage of children belonging to each subgroup. Scores in the low and medium subgroups remained stable, while scores in the high performance group increased by 11 points across age.

**Graph 3:**
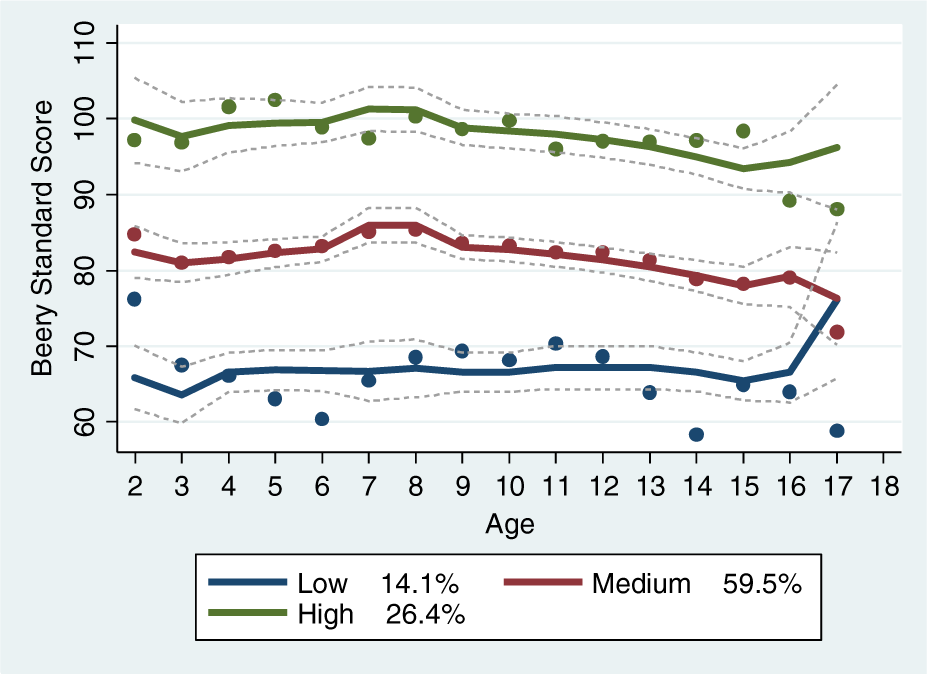
Beery Visuomotor Integration (VMI) Scale trajectory groups. This graph demonstrates Beery VMI trajectory subgroups characterized as high, medium and low performance and the percentage of children belonging to each subgroup. Scores of Beery VMI remained stable across age in all trajectory groups.

